# 4-1BB expression and signaling regulates MAIT cell activation and effector functions

**DOI:** 10.64898/2025.12.15.694466

**Authors:** R Lamichhane, J Williams, C Tirand, R Fouillé, LE Wedlock, A Poudel, RF Hannaway, JE Ussher

## Abstract

Mucosal associated invariant T (MAIT) cells are abundant unconventional T cells that recognize microbe-derived riboflavin metabolites presented by MR1. Upon recognition, they become activated, produce proinflammatory cytokines, chemokines and cytotoxic molecules. MAIT cells are also activated by cytokines independently of T cell receptor (TCR) engagement, however these two signals can also act in concert to finetune MAIT cell functions. Additionally, multiple other co-stimulatory signals have also been reported that can boost MAIT cell effector responses to TCR or cytokine stimulation. However, a comprehensive study exploring the role of surface bound TNF receptor superfamily (TNFRSF) molecule 4-1BB, well known for its co-stimulatory function during conventional T cell activation, is lacking in the context of MAIT cells. In this study, we show that 4-1BB is the earliest and most highly expressed TNFRSF co-stimulatory molecule on MAIT cells upon activation by Escherichia coli, with expression seen as early as six hours where it was predominantly MR1 mediated. 4-1BB expression on MAIT cells following late activation was due to both TCR signaling and cytokine signaling. We found marked differences in MAIT cell activation and cytokine expression between 4-1BB+ and 4-1BB- MAIT cells suggesting 4-1BB expression on MAIT cells is associated with functional superiority. By blocking 4-1BB signaling or co-culturing with a 4-1BBL overexpressing cell line we demonstrated an important role of co-stimulation via 4-1BB in MAIT cells during activation. Expression and signaling via 4-1BB also enhanced T-bet and Blimp1 expression. In summary, our study confirms a role for 4-1BB signaling during MAIT cell activation.

## Introduction

MAIT cells are abundant unconventional T cells in humans which express the semi-invariant T cell receptor (TCR) (TRAV1-2-TRAJ33/12/10 or Vα7.2-Jα33/12/20) and recognize pyrimidine derivatives from microbial riboflavin (vitamin B12) biosynthesis presented by MHC-I Related molecule (MR1) (1–3). In humans, MAIT cells consist of 1-10% of total T cells in blood and are enriched in mucosal tissues and liver, where they reach up to half of total T cells (4, 5). Multiple *in vitro* and *in vivo* studies have demonstrated their ability to recognize and be rapidly activated by cells infected with riboflavin-synthesising bacteria (6–9). TCR-independent activation can also occur by cytokines such as interleukin-12 (IL-12), IL-15 or IL-18 which are produced during bacterial and viral infection or inflammation (10–14). Activated MAIT cells produce various proinflammatory cytokines such as TNF-α, interferon-γ (IFNγ), and IL-17A, as well as cytotoxic molecules, including granzymes and perforin (15, 16). MAIT cells have been shown to protect against bacterial infection, as well as being involved in maintaining tissue homeostasis including tissue repair (9, 16–18). This makes MAIT cells one of the important immune players.

While strong TCR- or cytokine stimulation induces robust MAIT cell activation *in vitro*, a role for co-stimulation in the context of MAIT cells is also well demonstrated by multiple studies, both *in vitro* and *in vivo*. Co-stimulation during activation by a sub-optimal TCR or cytokine signal can rapidly boost the effector response of MAIT cells (14, 16, 19, 20). Most of the signals that have been reported to possess a co-stimulatory function in the context of MAIT cells are soluble cytokines, such as IL-6, IL-7, GM-CSF, IL-23, TNFα, TL1A and type I interferons (19–25). Studies investigating the role of co-stimulation via surface bound molecules are few and have mostly centered around CD8 co-stimulation. Gold et al. first reported the requirement of CD8 co-receptor engagement for enhanced IFNγ production by MAIT cells (defined as MR1-restricted *Mycobacterium tuberculosis*-reactive CD8^+^ T cell clones) by *M. tuberculosis*-infected dendritic cells (DCs) (26). Kurioka et al. demonstrated CD8 co-stimulation to have some role in MAIT cell activation and function in response to *Escherichia coli*-treated THP1 cells (27). Indeed, multiple studies have pointed out the functional superiority of CD8^+^ MAIT cells, which are also the dominant subset, over other CD8^−^ MAIT cell subsets (28–30). More recently, like the MHC-I molecule, Souter et al. highlighted a conserved CD8 binding site on MR1 and confirmed interaction between CD8 co-receptor and MR1 (31). A role for CD161 expression has also been demonstrated for cytokine but not cytotoxic molecule expression in MAIT cells activated by *E. coli*-infected HeLa cells (6). Few studies have examined the role of surface-bound co-stimulatory molecules which are activation induced (16). While MAIT cells have been shown to activate DCs both *in vitro* (32) and *in vivo* (33), and that this was partially dependent on CD40-CD40L interaction, this conferred no benefit to MAIT cell activation (32).

TNF receptor superfamily (TNFRSF) co-stimulatory molecules, such as 4-1BB (also known as TNFRSF9 or CD137), OX40 (also known as TNFRSF4 or CD134), CD30 (also known as TNFRSF8), CD27 (also known as TNFRSF7) and GITR (also known as TNFRSF18) are some of the most studied surface bound co-stimulatory molecules in the context of conventional T cells (34). Importantly, the inclusion of 4-1BB co-stimulation was shown to confer better survival with functional superiority and persistence of chimeric antigen receptor (CAR) T cells (35, 36). Although 4-1BB has been used as a readout of activation in a few studies, the significance of 4-1BB expression in the context of MAIT cells is only recently getting attention (32). Both TCR signals and cytokines, such as IL-12+IL-18 or IL-2, have been shown to upregulate 4-1BB expression on MAIT cells (16, 37). More recently 4-1BB expression was reported to be associated with higher IFNγ and IL-17A production by MAIT cells (37). Additionally, OX40 was recently found to regulate IL-9 production by MAIT cells from patients with gastritis (38). Interestingly, MAIT cells expressing 4-1BB or OX40 were highly proliferative (based on Ki-67 staining) compared to MAIT cells not expressing these molecules (37, 38). Therefore, it is important to understand the co-stimulatory function of these and other TNF superfamily members and how they regulate MAIT cell activation.

In this study, we demonstrate an important role of 4-1BB expression on MAIT cells during TCR mediated activation. Following activation, 4-1BB expressing MAIT cells possessed functional superiority with increased expression of T-bet and Blimp1. We further highlighted that MAIT cell function was modulated by 4-1BB ligation, suggesting 4-1BB as a potential target to finetune MAIT cell function.

## Results

### 4-1BB is expressed early on activated MAIT cells

To examine the expression of TNFRSF co-stimulatory molecules following MAIT cell activation, MAIT cells were activated for 6 hours with *E. coli* and the expression of 4-1BB, OX40 and CD30 on MAIT cells was assessed; 4-1BB (*TNFRSF9*), OX40 (*TNFRSF4*) and CD30 (*TNFRSF8*) were the top three significantly upregulated TNFRSF genes in MAIT cells activated by *E. coli* in our previously published transcriptomic study (30) (S. Figure 1A and B). *TNFRSF9* and *TNFRSF8* were also the most significantly upregulated TNFRSF genes in 5-A-RU and IL-12+IL-18 stimulated MAIT cells respectively (S. Figure 1C and D). As previously reported (16, 20), MAIT cells robustly expressed 4-1BB on the surface when activated by *E. coli* at 6 hours (Figure 1A-C) which is also consistent with the high expression of *TNFRSF9* in response to *E. coli* (Supplementary Figure 1A and B). By contrast, this *E. coli* mediated activation failed to induce detectable expression of OX40 and CD30 at 6 hours in comparison to the unstimulated control (Figure 1A-C), consistent with their lower transcript expression compared to *TNFRSF9* (S. Figure 1A and B). This suggests that 4-1BB is the earliest expressed TNFRSF co-stimulatory molecule following early *E. coli* mediated activation of MAIT cells.

**Figure 1:**
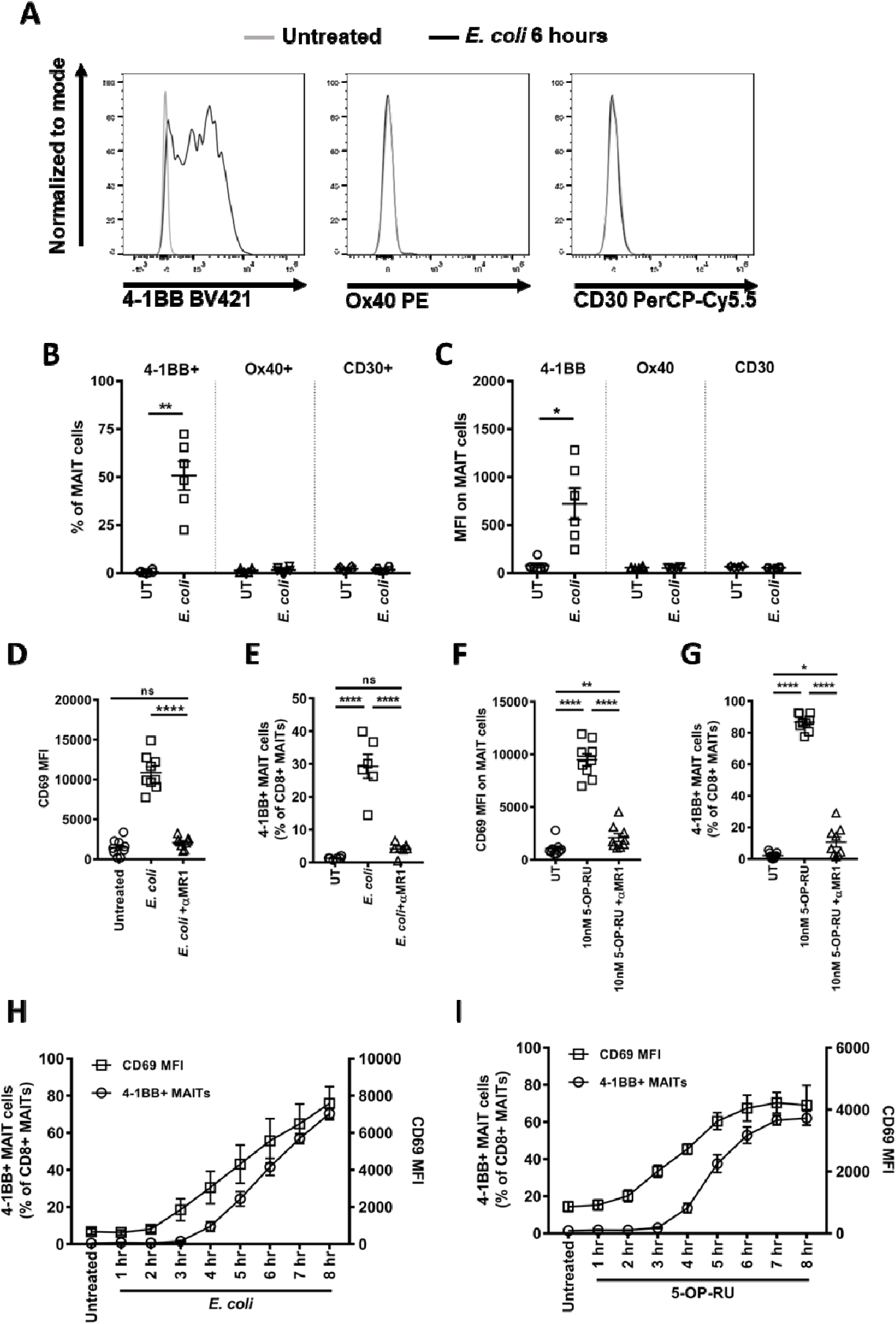
4-1BB is highly expressed on MAIT cells 6 hours after activation. (A-C) PBMCs were treated with *E. coli* (10 BpC) for 6 hours or left untreated and expression of 4-1BB, OX40 or CD30 was assessed on MAIT cells by flow cytometry. Representative flow cytometric plot (A), cumulative frequency (B) and mean fluorescent intensity (C) are shown. (D-I) PBMCs were treated with *E. coli* (10 BpC) ± anti-MR1 antibody (D and E) or 5-OP-RU ± anti-MR1 antibody (F and G) for 6 hours or left untreated and expression of 4-1BB or CD69 was assessed on MAIT cells by flow cytometry. (H and I) PBMCs were treated with *E. coli* (10 BpC) (H) or 10 nM 5-OP-RU (I) for different durations (1-8 hours) or left untreated. Data are presented as mean±SEM and are pooled from two independent experiments. Each symbol represents an individual healthy donor. Statistical analysis was performed using paired t-test (B and C) or one-way ANOVA with Sidak’s multiple comparison tests (D-G). ****p<0.0001, **p<0.01, *p<0.05, ns = non-significant, MFI = mean fluorescent intensity

Early activation of MAIT cells by riboflavin synthesizing bacteria is MR1 mediated (16). Consistent with this, early expression of both CD69 and 4-1BB on MAIT cells following activation by *E. coli* was significantly inhibited by the anti-MR1 antibody, confirming that the expression of 4-1BB at 6 hours is MR1 dependent (Figure 1D and E). MAIT cells also highly expressed 4-1BB when activated via their TCR by 5-OP-RU, the activating MR1 ligand (Figure 1F and G). The surface expression of 4-1BB on MAIT cells following stimulation with *E. coli* or 5-OP-RU was similar to that of CD69 and was dependent on the activation duration as well as the stimuli dose (Figure 1H and I and S. Figure 2). However, unlike CD69, which was already expressed at low levels on unstimulated MAIT cells, 4-1BB expression was only detectable upon activation.

**Figure 2:**
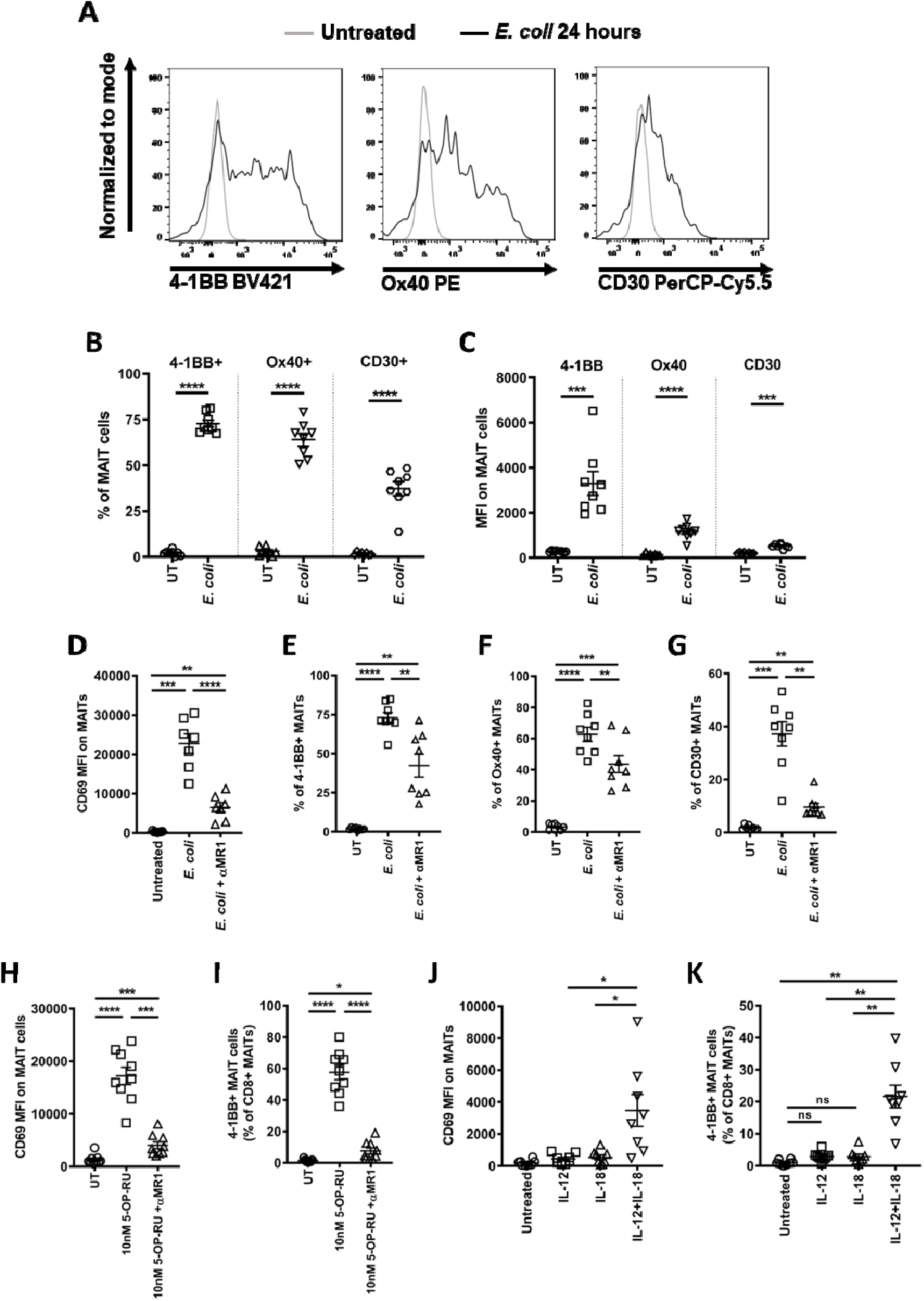
MAIT cells express multiple co-stimulatory molecules at 24 hours, mediated by both TCR and non-TCR signals. (A-C) PBMCs were treated with *E. coli* (10 BpC) for 24 hours or left untreated and expression of 4-1BB, OX40, and CD30 was assessed on MAIT cells by flow cytometry. Representative flow cytometric plots (A), cumulative frequency (B) and mean fluorescent intensity (C) are shown. (D-G) PBMCs were treated with *E. coli* (10 BpC) ± anti-MR1 antibody (D-G) for 24 hours or left untreated and expression of CD69 (D), 4-1BB (E), OX40 (F) and CD30 (G) were assessed on MAIT cells by flow cytometry. (H and I) PBMCs were treated with either IL-12, IL-18, or both (50 ng/mL each) for 24 hours and the expression of CD69 (H) and 4-1BB (I) was assessed on MAIT cells. (J and K) PBMCs were treated with 5-OP-RU (10 nM) ± anti-MR1 antibody for 24 hours or left untreated and expression of CD69 (J) and 4-1BB (K) were assessed on MAIT cells by flow cytometry. Data are presented as mean±SEM and are pooled from two independent experiments. Each symbol represents an individual healthy donor. Statistical analysis was performed using paired t-test (B and C) or one-way ANOVA with Sidak’s multiple comparison tests (D-K). ****p<0.0001, ***p<0.001, **p<0.01, *p<0.05, ns = non-significant, MFI = mean fluorescent intensity

Unlike *E. coli* stimulation for 6 hours, MAIT cells expressed all three TNFRSF co-stimulatory molecules (4-1BB, OX40 and CD30) following *E. coli* stimulation for 24 hours; more MAIT cells expressed 4-1BB and at higher levels (as measured by MFI) than OX40 or CD30 (Figure 2A-C). The expression of CD69, 4-1BB, OX40, and CD30 on MAIT cells stimulated by *E. coli* for 24 hours was partially inhibited upon MR1 blocking (Figure 2D-G). This suggests TCR independent signals, such as IL-12 and IL-18 which are produced in the PBMC system with *E. coli* treatment, also contribute to the expression of these TNFSF co-stimulatory molecules (12). Indeed, both IL-12 and IL-18 are required to significantly induce CD69 and 4-1BB expression on MAIT cells in the absence of TCR engagement (Figure 2H and I). Of note neither cytokine alone was able to significantly induce CD69 and 4-1BB expression, which is consistent with the previous observation that a combination of both cytokines is essential for robust MAIT cell activation (12). As at 6 hours, at 24 hours significant CD69 and 4-1BB expression on MAIT cells was seen in response to 5-OP-RU and was MR1-dependent (Figure 2J and K).

### 4-1BB+ MAIT cells are more activated and functionally superior

To understand the importance of 4-1BB expression on MAIT cells, we compared the expression of CD69 and production of TNFα and IFNγ by 4-1BB^+^ and 4-1BB^−^ MAIT cells following activation with *E. coli* for 24 hours; 4-1BB^+^ and 4-1BB^−^ MAIT cells were separated based on the untreated control as MAIT cells do not express 4-1BB in the absence of stimulation (Figure 1A and 3A). 4-1BB^+^ MAIT cells expressed significantly higher levels of CD69 (MFI) compared to 4-1BB^−^ MAIT cells (Figure 3B). The frequencies of TNFα or IFNγ producing MAIT cells were also significantly higher in the 4-1BB^+^ compartment compared to the 4-1BB^−^ compartment, suggesting 4-1BB expression is a marker for more functional MAIT cells (Figure 3C and D). CD69 expression was also higher on 4-1BB^+^ MAIT cells activated by *E. coli* at 6 hours (Figure 3E) or by cytokines (IL-12 and IL-18) at 24 hours (Figure 3F). Overall, 4-1BB^+^ MAIT cells were more activated with enhanced cytokine producing ability.

**Figure 3:**
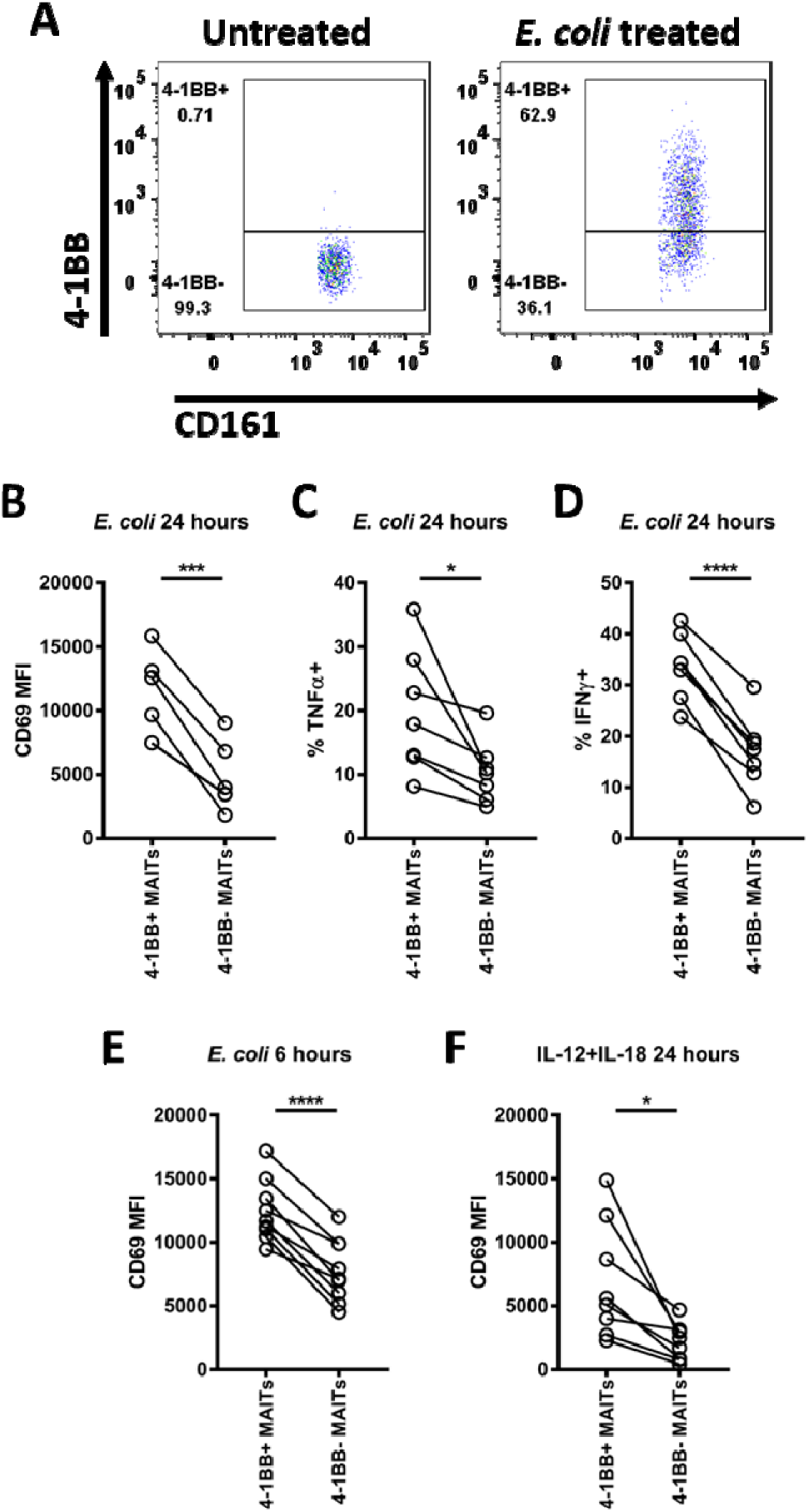
4-1BB expressing MAIT cells are more activated. (A-D) PBMCs were treated with *E. coli* (10 BpC) or left untreated for 24 hours and expression of CD69 (B), TNFα (C) and IFNγ (D) on 4-1BB^+^ and 4-1BB^−^ MAIT cells was compared by flow cytometry; unstimulated MAIT cells were used for gating to separate 4-1BB^+^ MAIT cells and 4-1BB^−^ MAIT cells (A). PBMCs were treated with *E. coli* (10 BpC) for 6 hours (E) or IL-12 and IL-18 (50 ng/mL each) for 24 hours (F) and expression of CD69 was assessed on MAIT cells. Data are presented as mean±SEM and are pooled from two independent experiments. Each symbol represents an individual health donor. Statistical analysis was performed using paired t-tests. ****p<0.0001, ***p<0.001, *p<0.05

### Signaling via 4-1BB is important for MAIT cell activation

To investigate if signaling via 4-1BB is important, we added anti-4-1BB antibody (clone BBK2) at different concentrations (1, 5 and 10 µg/mL) during MAIT cell activation by *E. coli* for 24 hours. A dose-dependent inhibition of the expression of CD69 and CD107a was observed following treatment with the 4-1BB antagonistic antibody; 10 µg/mL of anti-4-1BB antibody significantly reduced both CD69 and CD107a expression on MAIT cells (Figure 4A and B). Production of TNFα, IFNγ, and granzyme B by MAIT cells was also significantly reduced upon 4-1BB blocking, suggesting 4-1BB expression on MAIT cells positively regulates activation and the effector response during bacteria mediated MAIT cell activation (Figure 4C-F).

**Figure 4.**
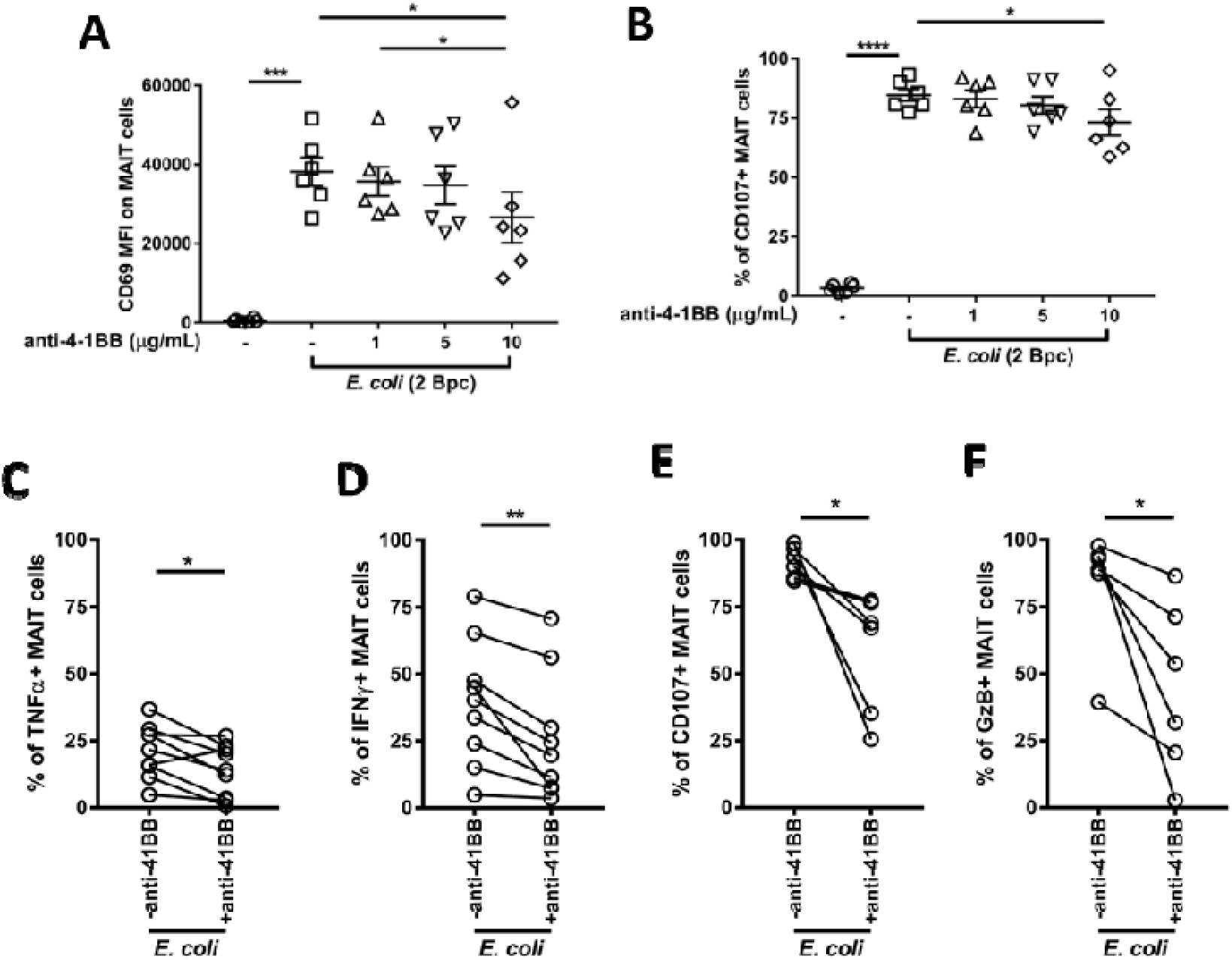
Signaling via 4-1BB is important for MAIT cell activation. (A and B) PBMCs were treated with *E. coli* (2 BpC) ± different concentrations of anti-4-1BB antibody (1, 5 and 10 µg/mL) for 24 hours and expression of CD69 (A) and CD107a (B) on MAIT cells was assessed by flow cytometry. (C-F) PBMCs were treated with *E. coli* (2 BpC) ± anti-4-1BB antibody (10 µg/mL) for 24 hours and expression of TNFα (C), IFNγ (D), CD107a (E) and granzyme B (F) was assessed on MAIT cells. Data are presented as mean±SEM and are pooled from two independent experiments. Each symbol represents an individual healthy donor. Statistical analysis was performed using one-way ANOVA with Sidak’s multiple comparison tests (A and B) or paired t-test (C-F). **p<0.01, *p<0.05.

### 4-1BB modulates the expression of T-bet and Blimp1 in MAIT cells

We recently showed that MAIT cells upregulate expression of multiple transcription factors upon activation, such as T-bet and Blimp1 (16). Consistently, expression of T-bet and Blimp1 were found to be upregulated in MAIT cells stimulated by *E. coli* for 24 hours (Figure 5A). Expression of these transcription factors is associated with better functional ability of MAIT cells (15, 16). Therefore, we compared the expression of T-bet and Blimp1 between 4-1BB^+^ and 4-1BB^−^ MAIT cells following *E. coli* stimulation to investigate whether functional superiority of 4-1BB^+^ MAIT cells was associated with enhanced expression of T-bet or Blimp1. Interestingly, expression of both T-bet and Blimp1 was higher in 4-1BB^+^ MAIT cells compared to 4-1BB^−^ MAIT cells, although only T-bet expression was significantly higher (5B and C). Next, we examined the expression of T-bet and Blimp1 following anti-4-1BB antibody treatment during *E. coli* mediated stimulation of MAIT cells to confirm a role for 4-1BB signaling in modulating T-bet and Blimp1 expression in MAIT cells. Expression of both T-bet and Blimp1 in MAIT cells was significantly diminished following 4-1BB blocking, confirming modulation of T-bet and Blimp1 expression in MAIT cells following 4-1BB signaling (5D and E).

**Figure 5.**
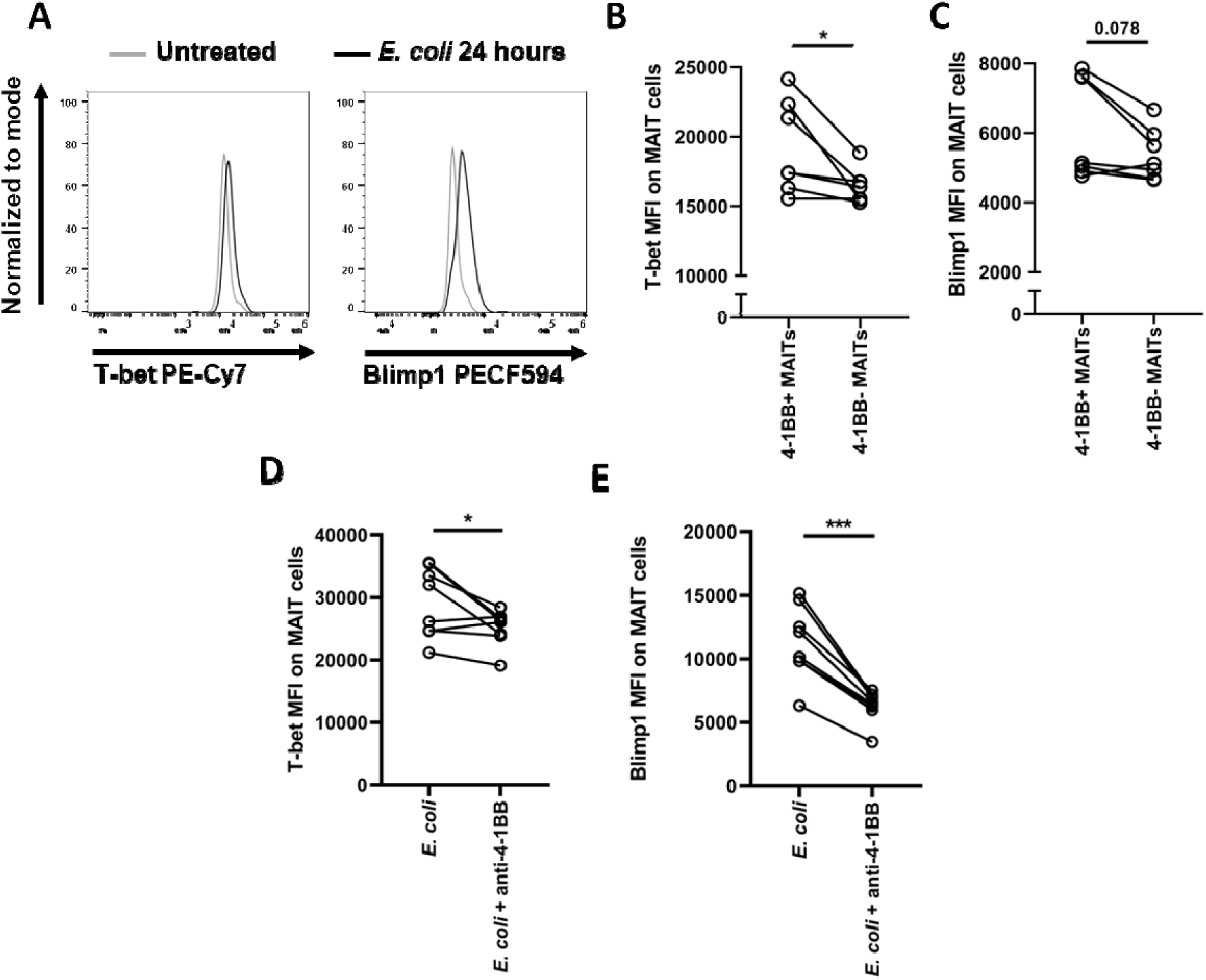
Expression of transcription factors are modulated upon 4-1BB signaling. (A) PBMCs were treated with *E. coli* (10 BpC) for 24 hours or left untreated and expression of T-bet and Blimp1 on MAIT cells was assessed by flow cytometry. (B and C) Expression of T-bet (B) and Blimp1 (C) on 4-1BB^+^ and 4-1BB^−^ MAIT cells was compared following treatment of PBMCs with *E. coli* (10 BpC) for 24 hours. (D and E) PBMCs were treated with *E. coli* (2 BpC) ± anti-4-1BB antibody (10 µg/mL) for 24 hours and expression of T-bet (D) and Blimp1 (E) was assessed on MAIT cells. Data are presented as mean±SEM and are pooled from two independent experiments. Each symbol represents an individual healthy donor. Statistical analysis was performed using paired t-test (B-E). ***p<0.001, *p<0.05. The p-value for near-significant comparison is shown.

### Enhanced 4-1BBL ligation augments MAIT cell activation and effector responses to TCR stimulation

4-1BB interacts with its ligand 4-1BBL which then initiates bidirectional co-stimulation (39). Therefore, we next sought to determine which cells expressed 4-1BBL in human PBMCs. Higher levels of 4-1BBL expression were detected on monocyte subpopulations compared to B and T cells, and expression was further increased with *E. coli* or IL-12+IL-18 treatment (S. Figure 3). Therefore, it seems a plausible hypothesis that provision of increased 4-1BBL expression and its ligation to 4-1BB enhances MAIT cell activation and function. To study this hypothesis, we sought to overexpress 4-1BBL on A549 cells (lung epithelial cells) as these cells are easy to transfect and have previously been shown to present bacteria-derived MR1 antigen to MAIT cells to activate them (8, 13). Consistent with this, when isolated CD8^+^ T cells were co-cultured with A549 cells and treated with different doses of 5-OP-RU, MAIT cells were activated in a dose-dependent manner, as determined by increased CD69 expression and downregulation of the Vα7.2 TCR expression (Figure 6A). Next, we generated a stable A549 cell line overexpressing 41BBL and confirmed 41BBL expression by flow cytometry (Figure 6B). Isolated CD8^+^ T cells were then co-cultured with either A549 parental cells or A549 cells overexpressing 4-1BBL and treated with 5-OP-RU or *E. coli* for 24 hours. MAIT cells co-cultured with 4-1BBL overexpressing A549 cells expressed significantly higher levels of CD69 and CD107a and produced significantly more TNFα and IFNγ in response to 10 nM 5-OP-RU compared with MAIT cells co-cultured with parental A549 cells (Figure 6C-F); although evident, these differences were less obvious with the lower dose (1 nM) of 5-OP-RU. Significant enhancement in TNFα and IFNγ production, but not CD69 or CD107a expression, was observed in the co-culture with 4-1BB overexpressing A549 cells when activated by *E. coli* (Figure 6C-F). Overall, this suggests that increased levels of 4-1BBL on antigen presenting cells enhances MAIT cell activation and their effector response to TCR stimulation.

**Figure 6:**
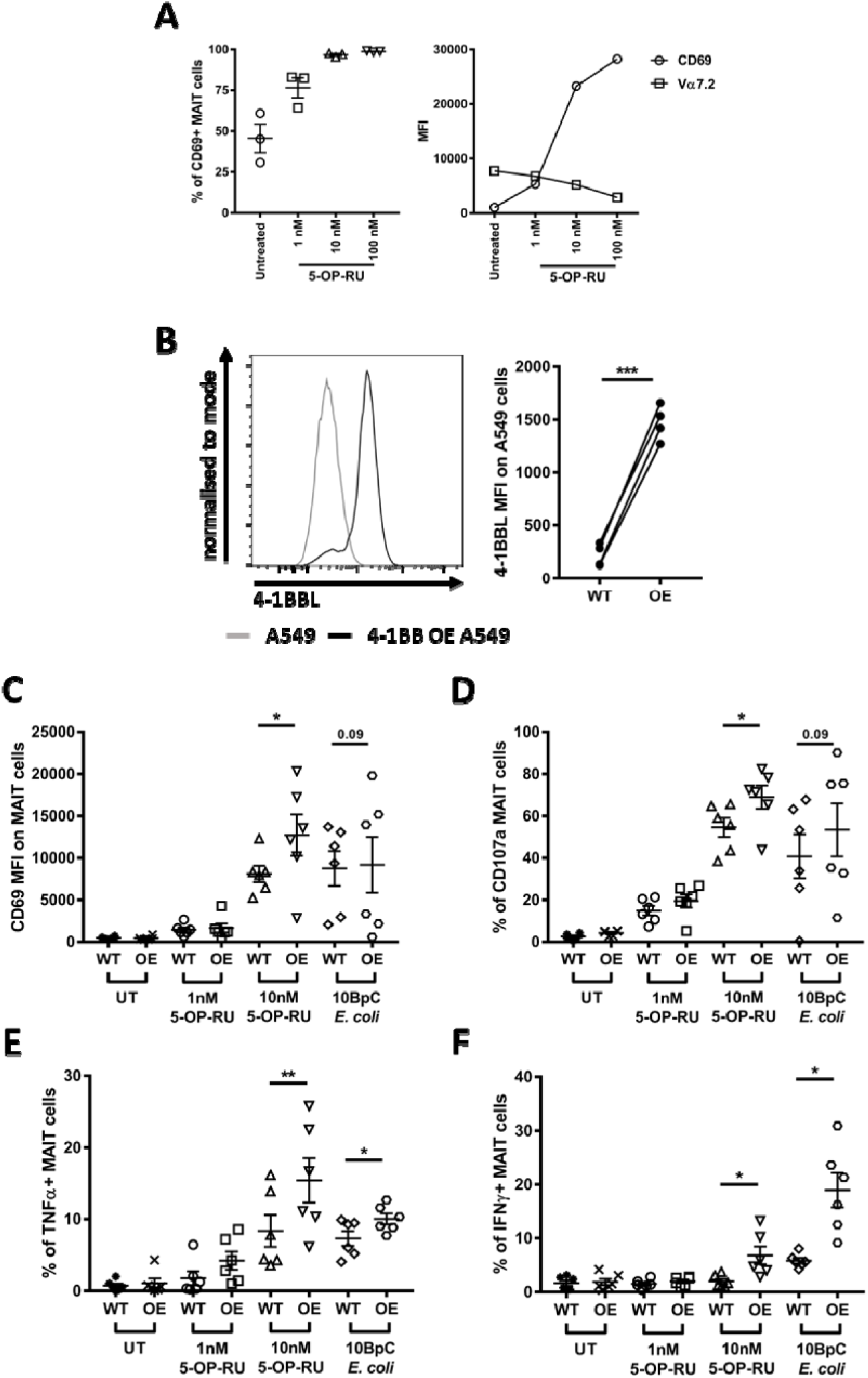
Late MAIT cell activation by 5-OP-RU or bacteria is enhanced by increased 4-1BBL expression. (A) Column enriched CD8^+^ T cells were co-cultured with A549 cells and treated with different concentrations of 5-OP-RU (1, 10, or 100 nM) for 24 hours and expression of CD69 and Vα7.2 TCR on MAIT cells was assessed by flow cytometry. (B) A549 cells were transfected with pcDNA3 containing human 4-1BBL and expression of 4-1BBL on transfected (over expressing, OE) and non-transfected (wild type, WT) A549 cells was compared by flow cytometry; representative flow cytometric plot and cumulative expression is shown from four independent experiments each performed in triplicate. (C-F) Column enriched CD8^+^ T cells were co-cultured with either non-transfected A549 cells or 4-1BBL overexpressing A549 cells and treated with either 1 or 10 nM 5-OP-RU or *E. coli* (10 BpC) for 24 hours and expression of CD69 (C), CD107a (D), TNFα (E) and IFNγ (F) was assessed on MAIT cells by flow cytometry. Data are presented as mean±SEM and are pooled from two (A, C-F) or four (B) independent experiments. Each symbol represents an individual healthy donor. Statistical analysis was performed using paired t-tests. **p<0.01, *p<0.05, MFI = mean fluorescent intensity. The p-value for near-significant comparisons is shown.

Next, we blocked TCR-MR1 interaction during activation by *E. coli* with anti-MR1 antibody or treated the co-culture with a combination of IL-12+IL-18 for 24 hours to examine the effect of 4-1BBL overexpression on MAIT cell activation by non-TCR signals (S. Figure 4). There was no enhancement in CD69 and CD107a expression and TNFα and IFNγ production by MAIT cells with TCR independent stimulation when co-cultured with 4-1BBL overexpressing A549 cells as opposed to A549 cells with basal 4-1BB expression (S. Figure 4A-D).

To further confirm that 4-1BBL ligation is important for TCR mediated MAIT cell activation, we treated the co-cultures with 5-OP-RU (10 nM or 100 nM) for 6 hours and assessed the expression of CD69 and CD107a and production of TNFα and IFNγ by MAIT cells (Figure 7). Co-culture with 4-1BBL overexpressing A549 cells significantly increased CD69 expression and TNFα production by MAIT cells (Figure 7A and C). A trend was also observed for increased CD107a expression by MAIT cells, but the change was not significant (Figure 7B); IFNγ expression was unchanged (Figure 7D). These observations confirmed that 4-1BB-4-1BBL interaction plays an important role in enhancing TCR-mediated MAIT cell activation and that this interaction can be targeted to finetune MAIT cell function.

**Figure 7:**
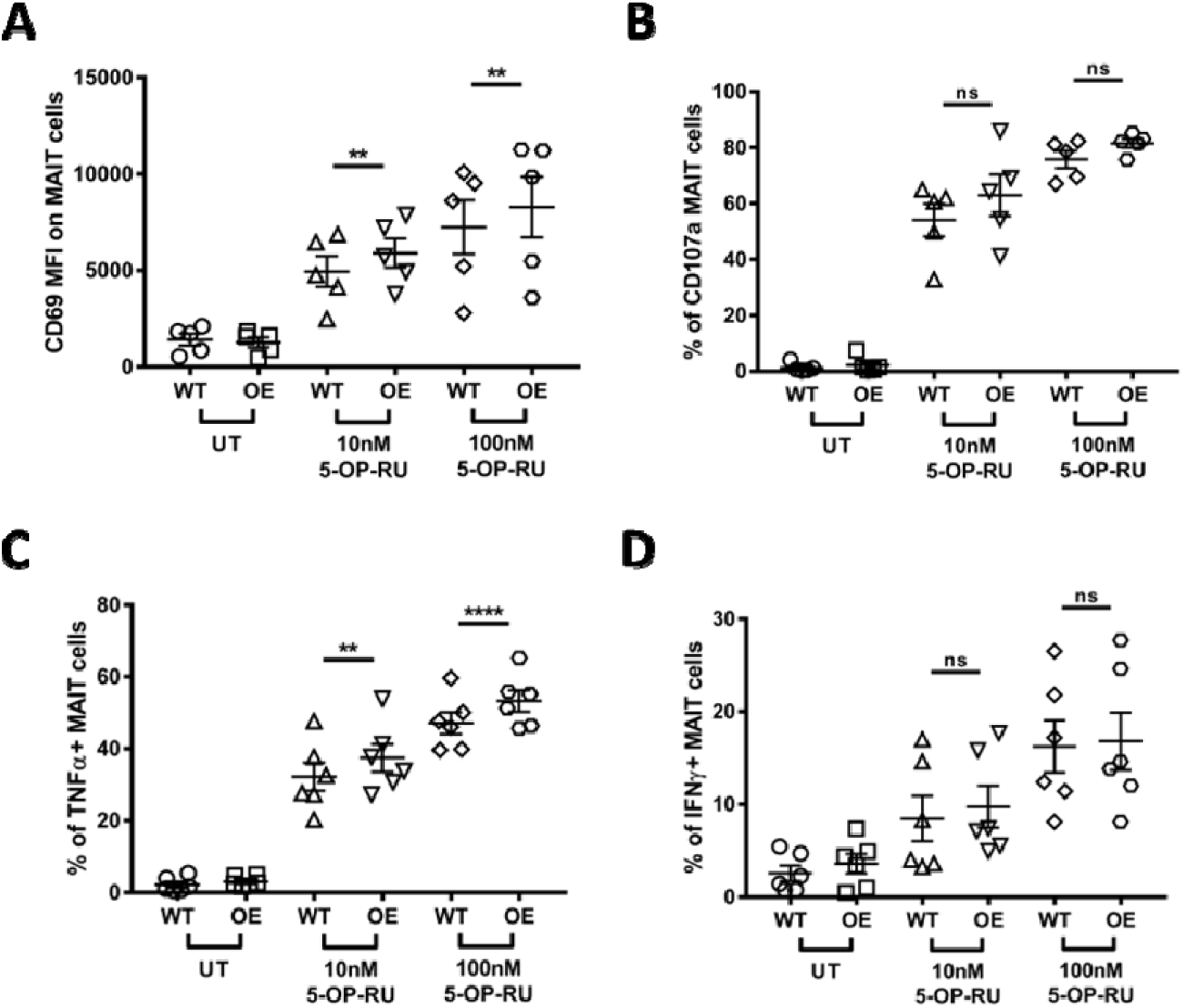
Early TCR-mediated MAIT cell activation is increased by increased 4-1BBL expression. Isolated CD8^+^ T cells were co-cultured with non-transfected A549 cells (wild type, WT) or 4-1BBL overexpressing (OE) A549 cells with either 10 or 100 nM 5-OP-RU for 6 hours and expression of CD69 (A), CD107a (B), TNFα (C) and IFNγ (D) was assessed on MAIT cells by flow cytometry. Data are presented as mean±SEM and are pooled from two independent experiments. Each symbol represents an individual healthy donor. Statistical analysis was performed using paired t-tests. ****p<0.0001, **p<0.01, ns = non-significant, MFI = mean fluorescent intensity

## Discussion

MAIT cells are an abundant unconventional T cell population in humans that respond to bacteria derived metabolites from the riboflavin synthesis pathway, and which are important in the control of bacterial infections. In this study, we have demonstrated an important role for the surface bound co-stimulatory molecule 4-1BB in regulating MAIT cell activation and effector functions in response to bacteria.

Expression of 4-1BB and its co-stimulatory role has been well characterized for conventional CD4^+^ and CD8^+^ T cells. The expression of 4-1BB is not limited to conventional T cells as multiple other immune cell populations, including NK cells and unconventional T cells including MAIT cells, iNKT cells, and γδ T cells are also reported to robustly express 4-1BB upon activation (16, 40–42) Consistent with this, we found that MAIT cells robustly express 4-1BB upon activation by bacteria (*E. coli*) or antigen alone (5-OP-RU) as early as 6 hours after stimulation. Interestingly, like mouse MAIT cells, (25) human MAIT cells did not express detectable 4-1BB in the absence of any stimulation; this was in contrast to CD69 expression, which a subset of MAIT cells expressed without stimulation, and therefore 4-1BB could serve as an alternative surface activation marker. Non-TCR signals also induced 4-1BB expression but to a lesser extent than TCR-mediated activation; non-TCR signals boosted 4-1BB expression on MAIT cells, acting in concert with the TCR signals during bacteria mediated activation. By contrast, more prolonged stimulation was needed for the expression of other members of TNFRSF family, OX40 and CD30, which was consistent with the published literature showing earlier expression of 4-1BB on murine CD8^+^ T cells (43).

4-1BB expression on T cells is well established to have a beneficial role during various infectious and non-infectious settings, including in cancer studies (37, 44, 45). In our study, MAIT cells expressing 4-1BB were more activated and possessed superior cytokine producing ability compared to MAIT cells that did not express 4-1BB following activation by bacteria (both early and late) or IL-12+IL-18. This was consistent with recent MAIT cell studies demonstrating the association of 4-1BB expression with enhanced IFNγ and IL-17 production by MAIT cells from blood as well as pleural effusions from patients with tuberculosis (37). Interestingly, in patients with tuberculosis, granzyme B expression was comparable between 4-1BB^+^ and 4-1BB^−^MAIT cells (37). Although granzyme B was not directly compared between 4-1BB^+^ and 4-1BB^−^MAIT cells in our study, when 4-1BB signaling was blocked with the antagonistic anti-4-1BB antibody (BBK4), reduced granzyme B expression was seen in MAIT cells, along with reduced activation (CD69), degranulation, and cytokine production, in response to *E. coli* treatment, suggesting signaling via 4-1BB is equally important for degranulation and cytotoxic functions. Expression of OX40, although not further assessed in this study, has been shown to result in better activation and enhanced IFNγ and IL-17 production by MAIT cells from diabetic patients (46). In addition to the well-established functions of MAIT cells, OX40 expression was also associated with IL-9 production by MAIT cells in *H. pylori* infection (47). A role of 4-1BB in inducing IL-13 production by murine CD4^+^ and CD8^+^ T cells (48) and a recent study highlighting that MAIT cells can be potent IL-13 producers (49) suggests the TNFRSF signaling might trigger unique effector response by MAIT cells which are still to be explored.

MAIT cells express a unique transcription factor profile that changes upon activation (16, 28, 29, 50). Consistent with previous studies, including ours, MAIT cells highly expressed T-bet and Blimp1 following activation by *E. coli* (15, 16). Enhanced expression of T-bet and Blimp1 has been associated with better activation and superior function, including enhanced IFNγ and granzyme B production by MAIT cells (15, 28). In this study, we observed higher expression of both T-bet and Blimp1 in 4-1BB^+^ MAIT cells than 4-1BB^−^ MAIT cells and significant reductions of both T-bet and Blimp1 in MAIT cells upon blocking 4-1BB signaling during *E. coli* mediated activation, suggesting that 4-1BB mediated co-stimulation modulates T-bet and Blimp1 expression. This is consistent with previous observations of significantly higher expression of IFNγ, granzyme B and T-bet in human CD8^+^ T cells following cis 4-1BB co-stimulation with respect to TCR stimulation (51) and higher *TBX21* and *PRDM1* transcript expression in CAR T cells against GD2-expressing solid tumors when the CD28 endodomain was replaced by the 4-1BB endodomain (44).

Activation also triggers 4-1BBL upregulation on multiple immune cells, including monocytes, the dominant antigen presenting cells in human PBMCs (41, 52). Testing the hypothesis that overexpressing 4-1BBL would increase co-stimulation and result in stronger T cell activation as reported previously in context of conventional T cells (53), we demonstrated that enhanced 4-1BBL expression on antigen presenting cells during TCR-mediated activation results in better activation and cytokine production by MAIT cells. In addition to increased activation and effector function, 4-1BB expressing MAIT cells have previously been shown to also express higher Ki-67, which was also observed following OX40 expression on MAIT cells (47). OX40 expression and signaling in MAIT cells has been associated with increased caspase-3 activation and activation induced cell death (46); whether the same is true of 4-1BB expression and signaling should be investigated. It may be that MAIT cells are more prone to apoptosis than conventional T cells following activation (54) and therefore apoptosis is not a signal specific phenomenon but merely fate of MAIT cells following activation. On the other hand, different co-stimulatory signals may result in different outcomes: replacing CD28 with 4-1BB endo-domain ameliorated exhaustion in CAR T cells during persistent antigenic stimulation (44). Given the benefits of 4-1BB expression and signaling, targeting 4-1BB might be useful to reinvigorate terminally exhausted tumour infiltrating MAIT cells (55, 56) or when employing MAIT cells for CAR T cell therapy; in recent years multiple studies have highlighted the potential anti-tumour function of MAIT cells, both *in vitro* and *in vivo* (57–59).

In summary, we have confirmed a co-stimulatory role for 4-1BB expression and signaling in MAIT cell activation and effector responses to TCR stimulation, which is mediated via the modulation of transcription factors, T-bet and Blimp1. Our study suggests that a deeper understanding of the regulatory mechanisms and full impact of 4-1BB signaling on MAIT cell activation, and their effect functions is warranted.

## Author contributions

JEU and RL designed the project and experiments. JEU managed the study. RL, JW, RF, CT, LW, RFH and AP acquired and analyzed data. RL and JEU conceived the work, interpreted the data and wrote the manuscript. All authors revised and approved the manuscript.

## Materials and methods

### Human peripheral blood mononuclear cells (PBMCs) isolation and cell culture

Blood was collected from healthy donors with prior informed written consent with the approval of the University of Otago Human Ethics Committee (H14/046). PBMCs were isolated by gradient centrifugation using Lymphoprep^TM^ (Axis-Shield, Oslo, Norway). Isolated cells were cryopreserved. Prior to experiments, cells were thawed and rested overnight in RPMI 1640 (Life Technologies) supplemented with 10% fetal bovine serum (FBS) (Gibco) and Penicillin-Streptomycin (Sigma-Aldrich). The same media was used for PBMC experiments. For some experiments, CD8^+^ cells were magnetically enriched using CD8 microbeads and MS columns following manufacturer’s instructions (Miltenyi Biotech).

### Bacteria, MR1 ligand, cytokines, and inhibitors

*Escherichia coli* was cultured overnight in Luria Bertani (LB) broth, was washed twice with PBS, and fixed with 2% paraformaldehyde for 20 mins. Bacteria were then washed, resuspended in PBS, and stored at 4 _C. Bacterial stocks were made fresh each month. Enumeration of *E. coli* was performed using 123 count eBeads (eBioscience, San Diego, USA) by flow cytometry.

5-A-RU was synthesized as previously described (16) and stored at −80 °C. 5-A-RU was mixed with methylglyoxal at 1:50 molar concentration and diluted with milliQ water as required for the experiments.

Anti-human MR1 antibody clone 26.5 (BioLegend) was used at 5 µg/mL. Human IL-12 (Miltenyi Biotech) and IL-18 (R & D Systems) were used at 50 ng/mL. Anti-human 4-1BB (Santa Cruz, clone BBK2) was used at concentrations indicated in the figure legend.

### A549 cell culture and transfection

DMEM supplemented with 10% FBS and Penicillin-Streptomycin was used for maintaining the lung epithelial cell line A549. The same media was used for all experiments involving co-culture of isolated CD8^+^ T cells with A549 cells.

A549 cells were transfected with the vector pcDNA3 containing the open reading frame of human 4-1BBL (40) using lipofectamine (LF) 2000. Briefly, cells were washed twice with warm PBS and left in plain DMEM until they were transfected with a mixture of OPTIMEM+pcDNA3 and OPTIMEM+LF2000 by gentle swirling. The cells were washed twice with PBS and maintained in DMEM supplemented with 10% FBS and Penicillin-Streptomycin and selection was performed with Geneticin (1 mg/mL).

### MAIT cell activation

A total of 1×10^6^ viable PBMCs/200µL were transferred to each of the wells in a U-bottom 96 well plate. *E. coli*, 5-OP-RU, or cytokines (IL-12 and IL-18, individually or in combination) at the concentrations indicated in figure legends were added at the start of the experiment. Except for the time course experiments, PBMCs were either treated for 6 hours with *E. coli* or 5-OP-RU, or 24 hours with the *E. coli* or the cytokines.

For the co-culture experiments, 1×10^5^ A549 (wild-type or 4-1BBL overexpressing) cells were seeded at the bottom of 48 well-plates and incubated overnight. The next day the media was changed and 1×10^5^ isolated CD8^+^ cells were added. Cells were either treated for 24 hours with 5-OP-RU, *E. coli*, or a combination of IL-12+IL-18, or for 6 hours with 5-OP-RU.

For blocking MR1-TCR or 4-1BB-4-1BBL interactions, 2.5 µg/mL anti-human MR1 antibody or 10 µg/mL anti-human 4-1BB antibody were added at the start of the experiments respectively. For some experiments examining intracellular proteins in MAIT cells (TNFα, IFNγ and granzyme B), brefeldin A (BioLegend) was added at 5 µg/mL for the final four hours of incubation. CD107a PE was added at the start of the experiment to capture any CD107a coming to the cell surface during the incubation period.

### Immunostaining and flow cytometry

At the end of the treatment, PBMCs were washed with PBS and stained for the surface proteins (CD69, 4-1BB, OX40, CD30 and CD161 for MAIT cells, or CD14, CD16, CD20 and 4-1BBL for monocytes and B cells) for 25 mins, followed by washing with PBS and fixing with 2% paraformaldehyde for 20 mins. Cells were washed with PBS, permeabilized with Perm wash buffer (BioLegend) and stained (CD3, CD8, Vα7.2, TNFα, IFNγ, granzyme B) for 25 mins. This was followed by two washes with Perm wash buffer and two washes with PBS. Transcription factor staining was performed using eBioscience™ FOXp3 / Transcription Factor Staining Buffer Set (Invitrogen) following manufacturer’s instructions. Samples were acquired on a FACSCanto^TM^ II or LSRFortessa^TM^ (both from BD) and analyzed by FlowJo^TM^ V10 (TreeStar, Ashland, USA). Antibodies used were: CD3 PE-Cy7 (UCHT1) or BV510 (OKT3), Vα7.2 TCR PE or PE-Cy7 or AF700 (3C10), CD161 APC or BV605 (HP-3G10), CD107a PE (H4A3), CD69 FITC (FN50), TNF-α FITC (Mab11), IFN-γ PerCP-Cy5.5 (4S.B3), granzyme B FITC (QA16AO2), 4-1BB PE or BV421 (4B41), OX40 PE (Ber-ACT35), CD30 PE-Cy7 (BY88), 4-1BBL APC (5F4), CD14 FITC (S18004C), CD16 PE (3G8), CD20 APC-Cy7 (2H7), T-bet PE-Cy7 (4B10), (all from BioLegend), CD8 eFluor450 (RPA-T8, eBioscience) and Blimp1 PECF594 (6D3, BD Biosciences).

### Statistical analysis

All data are presented as mean ±SEM and were analysed in GraphPad Prism V8.4. Data were tested for normality before any statistical comparison. Two-tailed paired t-test or one-way ANOVA with Sidak’s multiple comparison test was used for the statistical comparisons between two groups and between multiple groups respectively. A p-value of 0.05 or less was considered significant.

## Acknowledgement

We thank Dr Andrea Vernall (Department of Chemistry, University of Otago, New Zealand) for the synthesis and supply of 5-A-RU. The vector pcDNA3 with the human 4-1BBL was a kind gift from Professor Helmut Rainer Salih, Eberhard-Karls University, Tuebingen, Germany.

---

**S. Figure 1:**
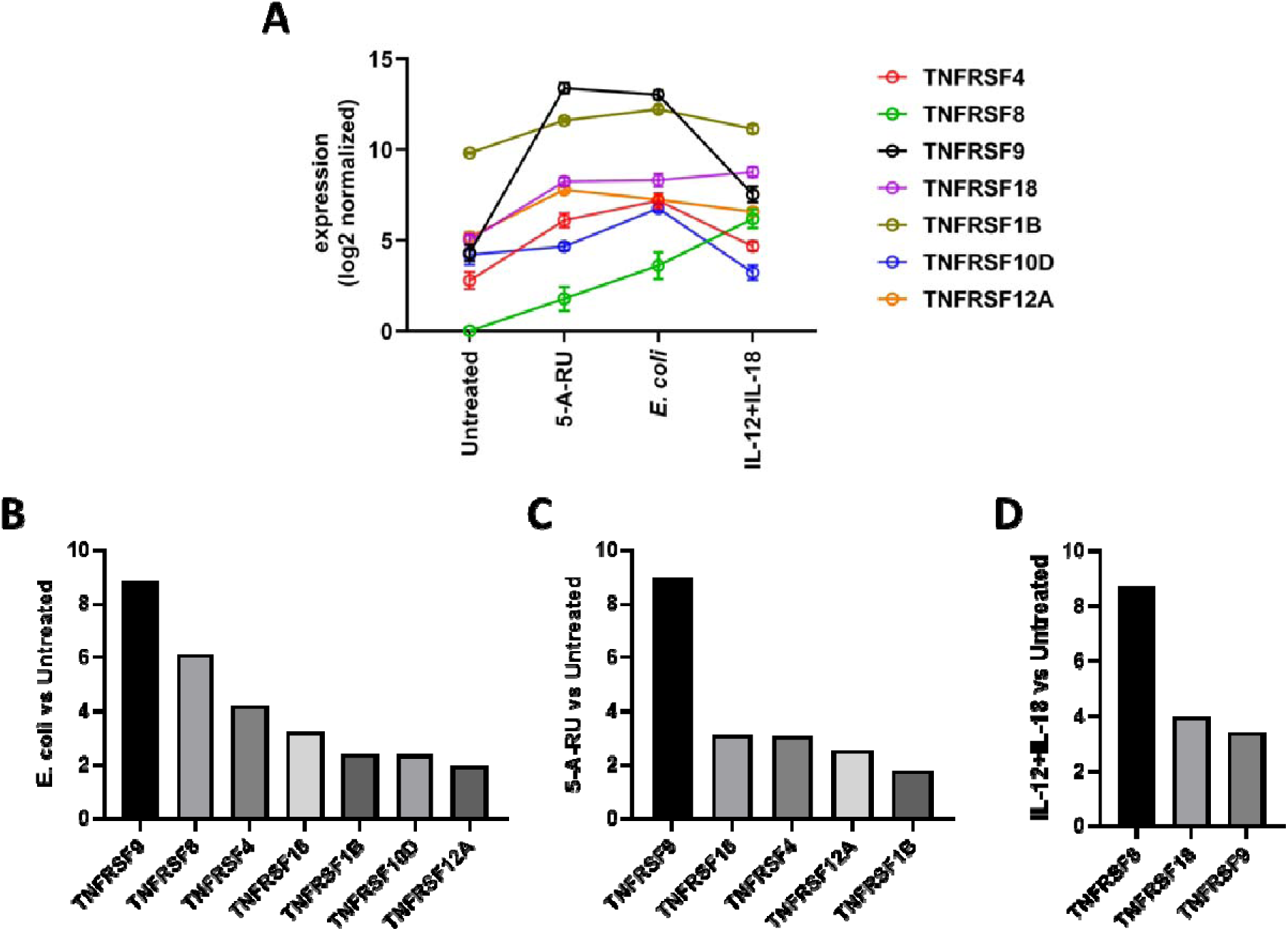
Expression of TNFRSF co-stimulatory genes in MAIT cells following activation. TNFRSF9 is highly expressed TNFRSF co-stimulatory gene in MAIT cells upon activation by TCR signals. Expression of differentially expressed TNFRSF genes in MAIT cells against either of the three different stimuli (E. coli or 5-A-RU for 6 hours, or IL-12+IL-18 for 24 hours) vs Untreated control (A) and fold change value of the genes in MAIT cells that are significantly differentially expressed by each treatment compared to Untreated (B-D). Data are presented as mean±SEM (n = 7) (A).

**S. Figure 2:**
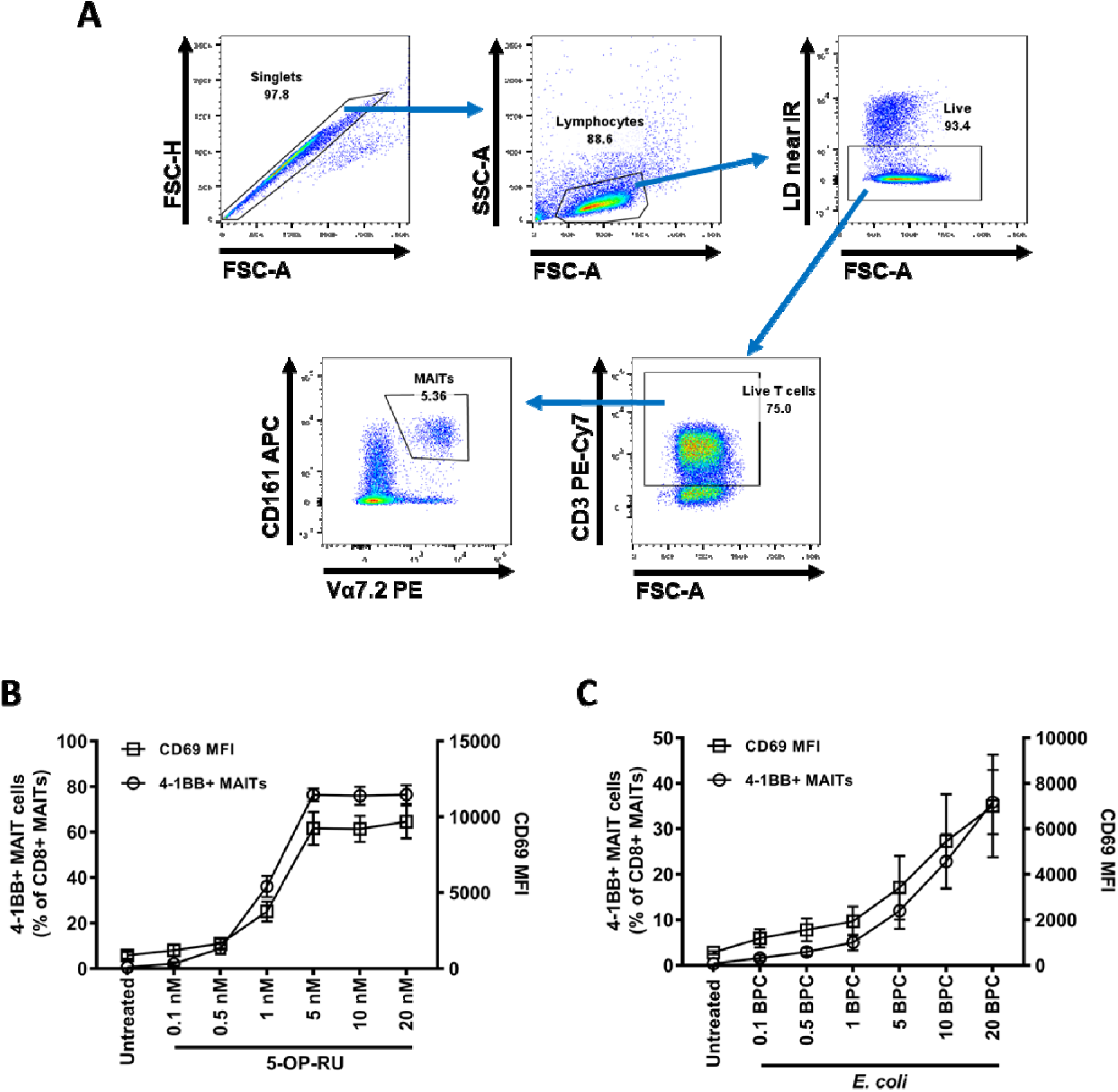
Similar modulation of 4-1BB and CD69 expression on MAIT cells by different doses of 5-OP-RU and *E. coli* at 6 hours. PBMCs were treated with different concentrations of 5-OP-RU (0.1, 0.5, 1, 5, 10, or 20 nM) (B) or *E. coli* (0.1, 0.5, 1, 5, or 20 BpC) (C) for 6 hours or left untreated. Expression of 4-1BB (% of cells expressing) or CD69 (MFI) on MAIT cells was assessed by flow cytometry; gating for identification of MAIT cells is shown (A). Data are presented as mean±SEM and are pooled from two independent experiments.

**S. Figure 3:**
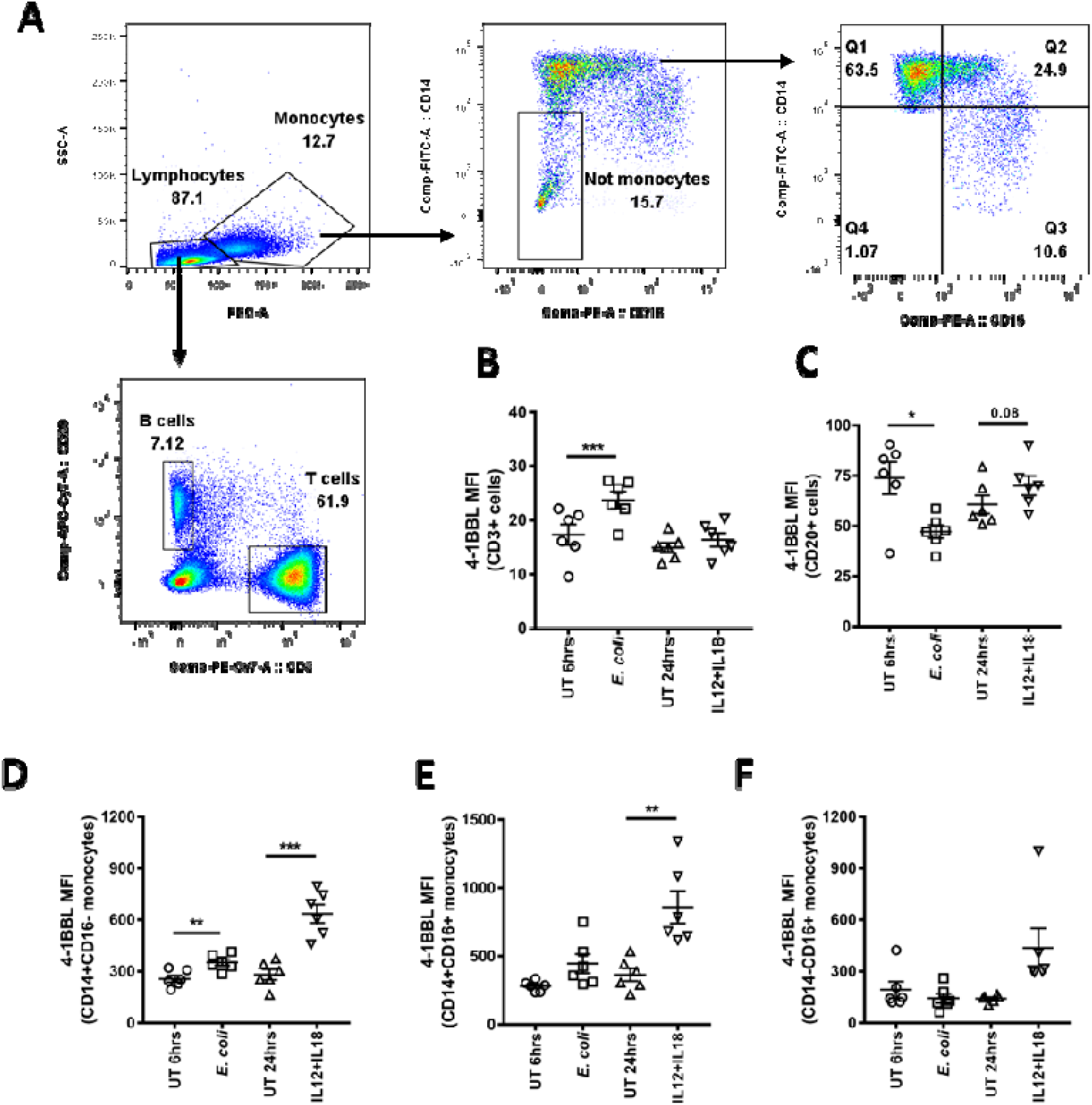
Primary human cells express 4-1BBL. PBMCs were treated with *E. coli* (10 BpC) for 6 hours or IL-12+IL-18 (50 ng/mL each) for 24 hours or left untreated. Expression of 4-1BBL on T cells (B), B cells (C) and different monocyte subsets (D-F) was assessed by flow cytometry; gating strategy from live single cells is shown (A). Data are presented as mean±SEM and are pooled from two independent experiments. Each symbol represents individual healthy donor. Statistical analysis was performed using paired t-tests. ***p<0.001, **p<0.01, *p<0.05. The p-value for near-significant comparison is shown.

**S. Figure 4:**
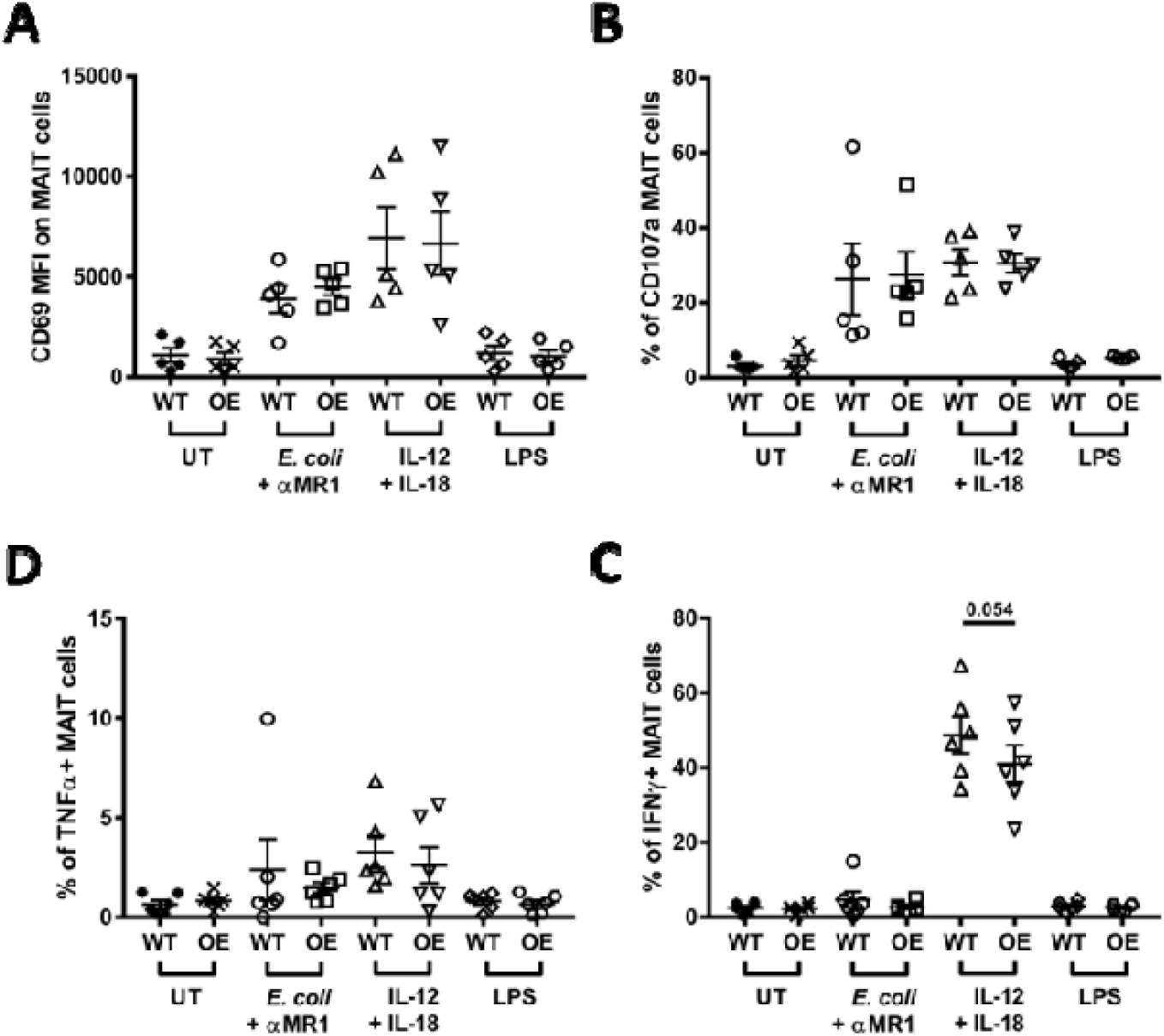
MAIT cell activation by cytokines is unaffected following increased 4-1BBL expression. Column enriched CD8^+^ T cells were co-cultured with non-transfected A549 cells (wild type, WT) or 4-1BBL overexpressing (OE) A549 cells with either *E. coli* (10 BpC) + anti-MR1 antibody, the combination of IL-12+IL-18 (50 ng/mL each), or LPS (1 μg/mL) for 24 hours and expression of CD69 (A), CD107a (B), TNFα (C), and IFNγ (D) was assessed on MAIT cells by flow cytometry. Data are presented as mean±SEM and are pooled from two independent experiments. Each symbol represents an individual healthy donor. Statistical analysis was performed using paired t-tests. *p<0.05. The p-value for near-significant comparison is shown. MFI = mean fluorescent intensity

